# A tRNA-acetylating toxin and detoxifying enzyme in *Mycobacterium tuberculosis*

**DOI:** 10.1101/2021.12.20.473312

**Authors:** Francesca G. Tomasi, Alexander M. J. Hall, Jessica T. P. Schweber, Charles L. Dulberger, Kerry McGowen, Qingyun Liu, Sarah M. Fortune, Sophie Helaine, Eric J. Rubin

## Abstract

Toxin-antitoxin (TA) systems allow bacteria to adapt to changing environments without altering gene expression. Despite being overrepresented in *Mycobacterium tuberculosis* (*Mtb*), their individual physiological roles remain elusive. We describe a TA system in *Mtb* which we have named TacAT due to its homology to previously discovered systems in Salmonella. The toxin, TacT, blocks growth by acetylating glycyl-tRNAs and inhibiting translation. Its effects are reversed by the enzyme peptidyl tRNA hydrolase (Pth), which also cleaves peptidyl tRNAs that are prematurely released from stalled ribosomes. Pth is essential in most bacteria and thereby has been proposed as a promising drug target for complex pathogens like *Mtb*. Transposon sequencing data suggest that the *tacAT* operon is nonessential for *Mtb* growth *in vitro*, and premature stop mutations in this TA system present in some clinical isolates suggest that it is also dispensable *in vivo*. We assessed whether TacT modulates *pth* essentiality in *Mtb*, as drugs targeting Pth might be ineffective if TacAT is disrupted. We find that *pth* essentiality is unaffected by the absence of *tacAT*. These results highlight a fundamental aspect of mycobacterial biology and indicate that Pth’s essential role hinges on its peptidyl-tRNA hydrolase activity. Our work underscores Pth’s potential as a viable target for new antibiotics.

## Introduction

*Mycobacterium tuberculosis* (*Mtb*), which causes tuberculosis (TB), is a leading cause of global infectious disease mortality[1]. *Mtb*’s ability to regulate its growth in different stressful conditions *in vitro* is thought to be an important part of its success *in vivo*. One of this pathogen’s tools for growth regulation is an expansive network of toxin antitoxin (TA) systems, with at least 100 putative modules that encompass nearly 4% of *Mtb*’s coding capacity[2, 3]. Most of these systems in *Mtb* can be grouped into five main mechanistic families based on sequence homology: VapBC, MazEF, RelBE, HigBA, and ParDE[4, 5].Toxins of TAs are characterized by their general intracellular targets and mechanisms of activity with most known *Mtb* toxins being RNAses that cleave rRNA, mRNA, or tRNA.

Most *Mtb* toxins are classified as Type II toxin-antitoxins, the most widespread and heavily studied type. In these systems, a protein antitoxin is bound tightly to its cognate protein toxin and acts to neutralize it[6]. If the antitoxin is degraded, the toxin assumes its active form and blocks an essential process such as DNA replication or protein synthesis until antitoxin production resumes[6]. Despite being widespread in bacteria, the physiological roles of TA systems are just emerging, with some having been linked to plasmid maintenance, bacteriophage immunity, and the formation of dormant, antibiotic-tolerant persisters[7, 8]. TA systems might play a role in *Mtb*’s ability to withstand host and antibiotic pressures by controlling growth under different stress conditions[4, 9, 10]. However, it remains to be determined whether or to what extent they play a role in pathogenesis. A significant barrier to understanding TA systems is the challenge of directly measuring native toxin activity in cells, and therefore understanding when they are active and how they interact with other enzymes. Because of the nature of TA system autoregulation and post-translational control, transcription upregulation data alone do not necessarily indicate toxin activation[11]. So far, studies investigating TA systems in bacteria often measure activity in cells using ectopic overexpression constructs[4, 5, 12]. These studies offer fascinating mechanistic insights but do so in isolation from other intracellular systems, and it has been difficult to link the molecular mechanisms of TA systems to their biological roles.

Recently, a new class of TA systems called TacAT was discovered in Salmonella and homologs have since been identified in other species including *Escherichia coli* and *Klebsiella pneumoniae*[13–17]. The TacT toxins in this family are GCN5-related N-acetyltransferases that acetylate aminoacyl tRNA and block incorporation of an amino acid into a growing peptide chain. TacT’s unique mechanism of action – which can be detected using liquid chromatography-coupled mass spectrometry – makes it an appealing TA system to study in the context of bacterial physiology[13, 14, 18, 19]. An unusual aspect of TacAT systems is that, while the antitoxin can block toxin activity as seen with other TA systems, the effect can also be reversed via the ubiquitous and essential bacterial enzyme peptidyl tRNA hydrolase (Pth)[13, 14]. This enzyme cleaves acetylated amino acids from tRNA molecules, effectively unblocking protein synthesis. TacT’s mechanistic connection to an essential enzyme makes it an appealing TA system to study in the context of gene essentiality.

Here we describe an *Mtb* homologue of the TacAT TA system, the first of its kind to be identified in this organism. We show that this TA system is encoded by the Rv0918-0919 operon and confirm that Rv0919 encodes a tRNA-acetylating toxin whose activity can be reversed by *Mtb* Pth (Rv1014c). While *pth* is required for growth in *Mtb*, transposon sequencing data suggest that the *Mtb* TacAT operon is dispensable for growth *in vitro*[20]. We have also identified premature stop mutations in this TA system in clinical isolates, suggesting it is not under positive selective pressure clinically. If TacT activity modulates *pth* essentiality in *Mtb*, then drugs targeting Pth might be ineffective if TacAT activity is disrupted, as has already happened in clinical isolates. However, we find that while the *tacAT* operon is indeed dispensable, *pth* essentiality is is not, and its requirement for *Mtb* growth is unaffected by the absence of this TA system. Our results indicate that Pth’s essential role in *Mtb* hinges on its function in cleaving peptidyl-tRNA and not acetylated aminoacyl tRNA. Our work underscores Pth’s potential as a viable target for new antibiotics, while also highlighting multiple angles from which to study TA systems in *Mtb*.

## Results

### Rv0918-0919 encodes a toxin-antitoxin system that inhibits growth by acetylating glycyl-tRNAs

Previous studies have identified over 100 putative TA systems in *Mtb*, based on genetic architecture and homology to known TA systems[5]. The *Mtb* operon Rv0918-0919 has been computationally flagged as a possible TA system due to its polycistronic organization and the presence of a conserved, DNA-binding RHH domain in the putative antitoxin gene, Rv0918 (Figure 1A)[5, 21]. Rv0919 contains a conserved GNAT domain, and protein BLAST results show >50% sequence identity to the N-acetyltransferase TacAT toxins in *Salmonella* (Figure 1A)[22].

**Figure 1:**
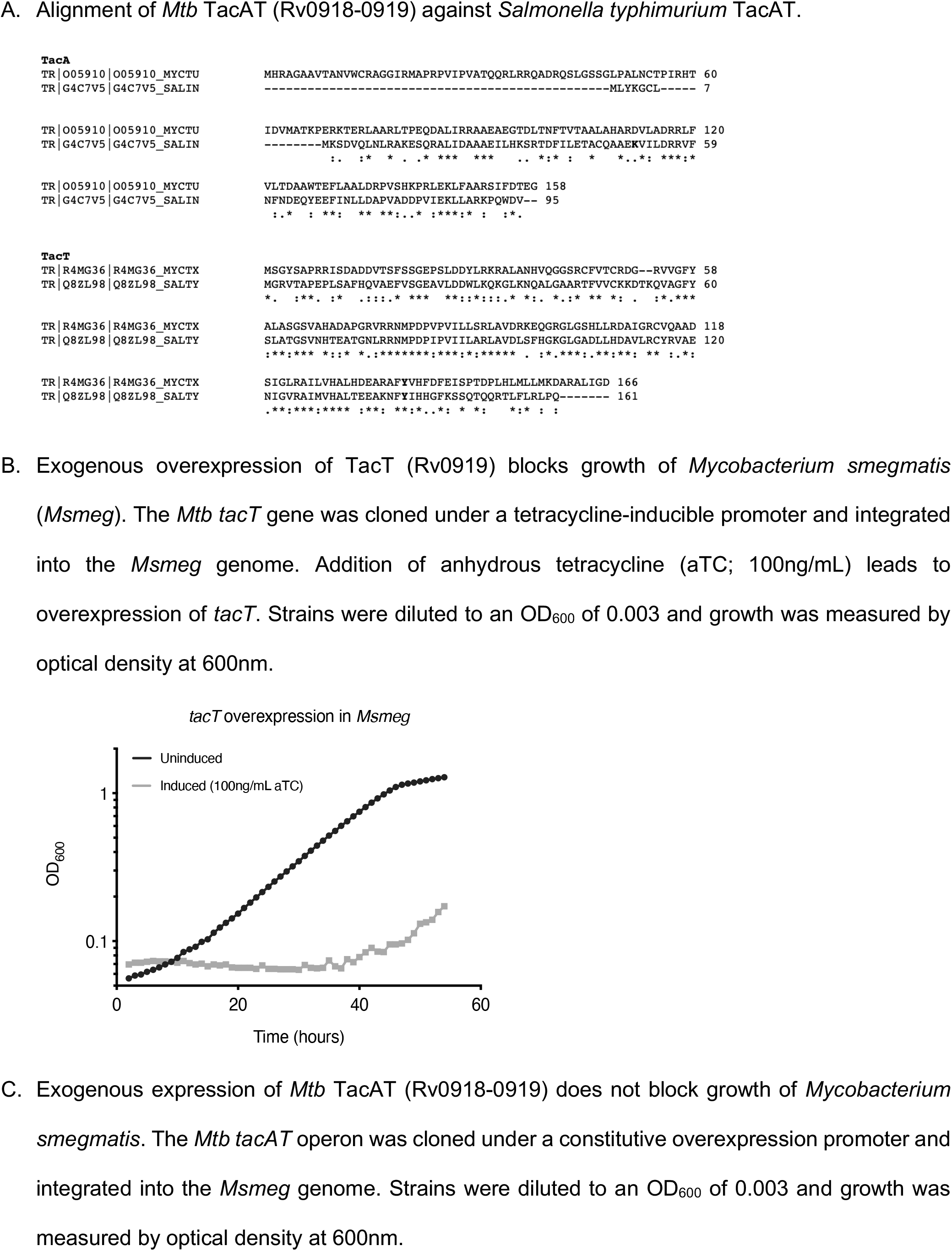
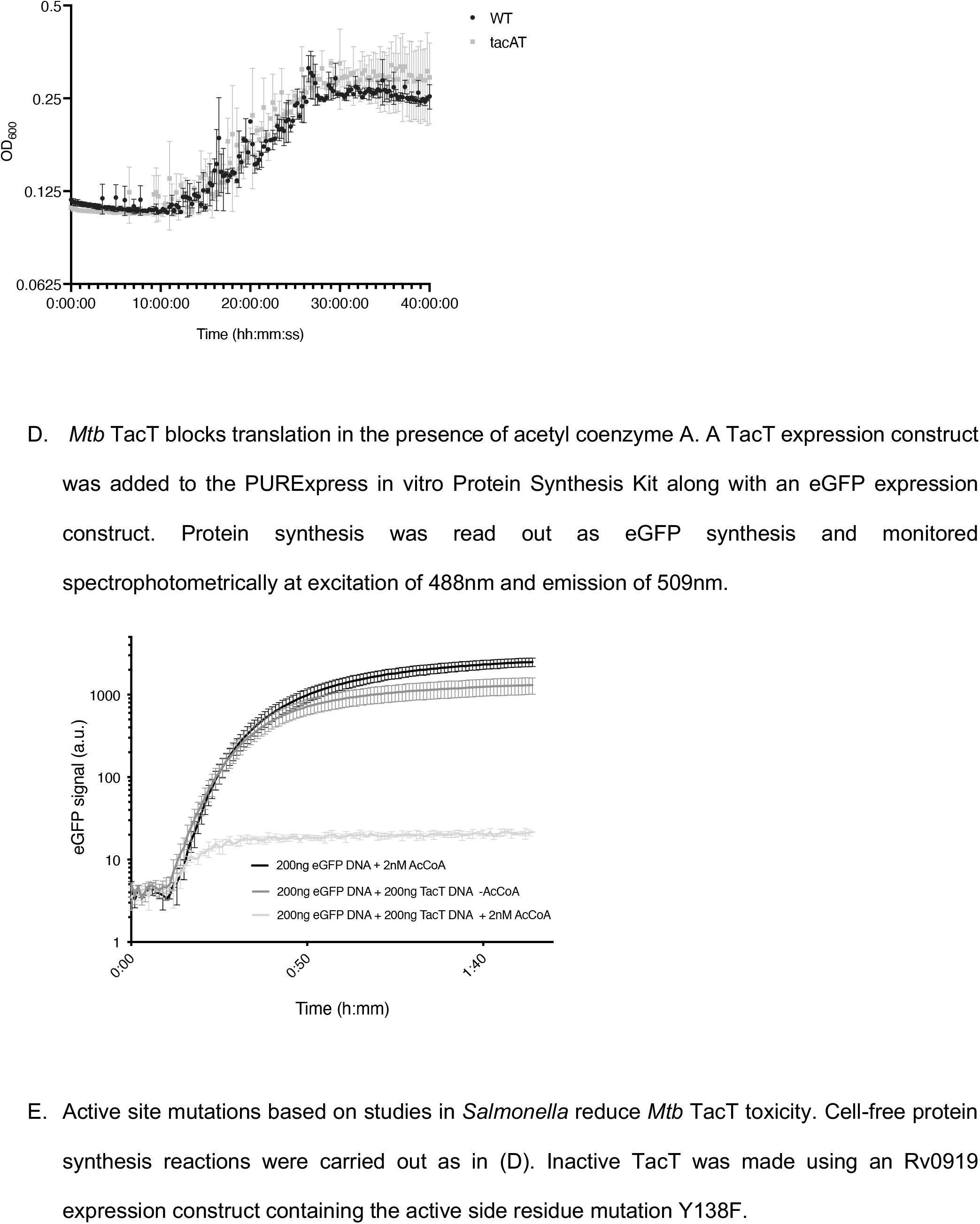
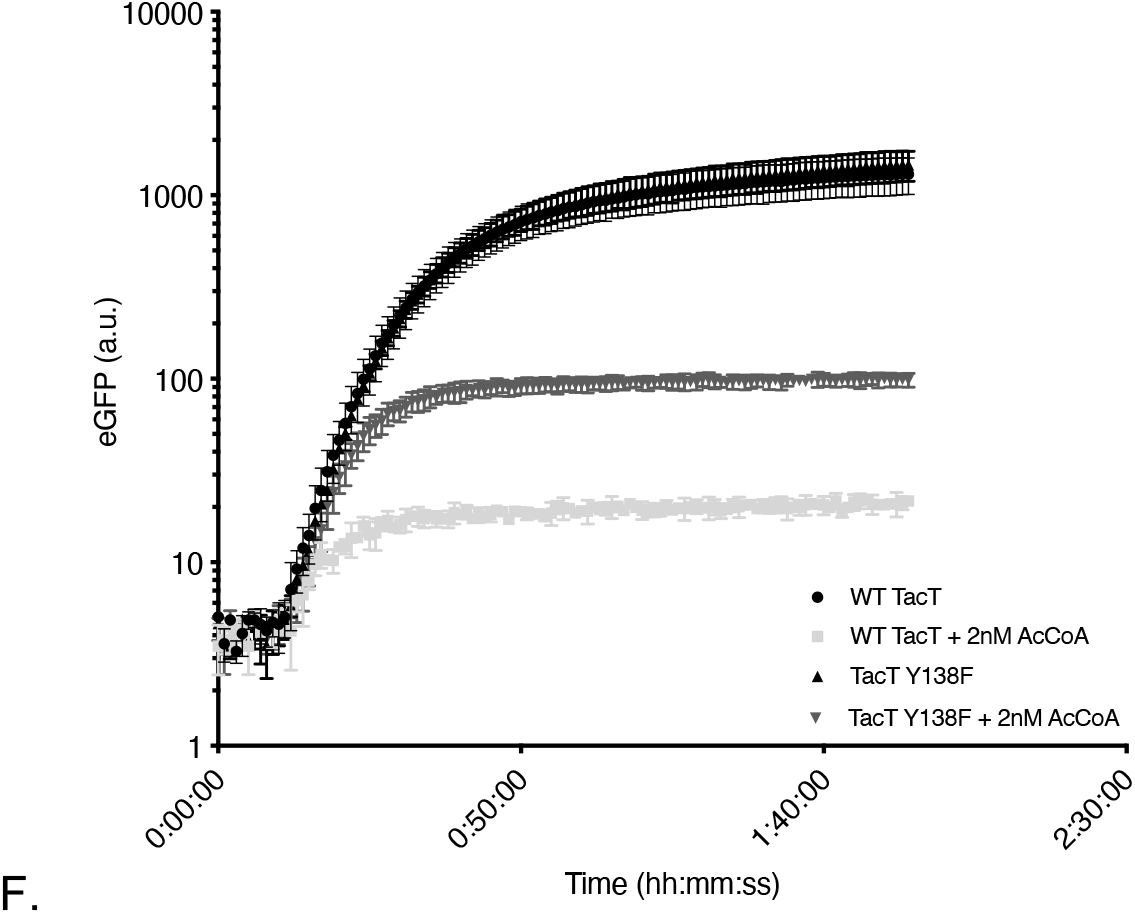
TacT is a toxin that inhibits protein synthesis.

We hypothesized that Rv0919 encodes a TacT-like toxin that inhibits growth by blocking translation. The closely related but faster-growing, non-pathogenic model organism *Mycobacterium smegmatis* (*Msmeg*) does not encode any putative TacAT-like systems[21, 22]. Therefore, to study *Mtb* TacT in isolation from other potential interacting genes we built an integrating vector carrying Rv0919 under the control of an anhydrous tetracycline (aTC)-inducible promoter. Induced overexpression of Rv0919 in *Msmeg* inhibited growth (Figure 1B), while constitutive expression of the entire Rv0918-0919 operon did not (Figure 1C), showing that Rv0919 encodes a growth-inhibiting enzyme that is not active in the presence of Rv0918.

We next assessed Rv0919 activity *in vitro* using a cell-free protein synthesis kit. We found that, while GFP could be efficiently expressed in this system, adding a DNA construct encoding Rv0919 blocked synthesis, though only when acetyl co-enzyme A was added (Figure 1D). This suggests that Rv0919 uses acetyl co-enzyme A as an acetyl group donor, as has been seen with other TacAT systems[14]. A construct encoding Rv0919 with an active site mutation homologous to one identified in Salmonella (Y138F in *Mtb*) only partially abrogated protein synthesis, while Rv0919 with two different predicted catalytic site mutations (A91P and Y138F) had no ability to inhibit GFP synthesis, even in the presence of acetyl co-enzyme A (Figure 1E)[13]. These results suggest that Rv0919 inhibits growth by acetylating a component of the protein synthesis apparatus.

All other described TacAT-like systems encode a toxin that acetylates the amino acid on charged tRNA. Different organism toxins acetylate different tRNAs. For instance, in *Salmonella*, three different TacT-like toxins have been described. These block elongation by acetylating glycyl, isoleucyl-, leucyl-, and, to a lesser extent, other aminoacyl tRNAs *in vitro* [13] but solely glycyl-tRNA *in vivo* (in preparation). Meanwhile, in *E. coli*, the GNAT toxin AtaT was initially thought to block initiation of protein synthesis by acetylating methionine on initiator fMet-tRNA [17] but was more recently reported to acetylate preferentially glycyl-tRNA alongside others[19]. We hypothesized that *Mtb* TacT also acetylates charged tRNAs. To identify if any and which tRNA species might be affected by this enzyme, we purified total RNA from *Msmeg* overexpressing *Mtb* TacT and used liquid chromatography-coupled mass spectrometry (LCMS) to analyze tRNA acetylation (Figure 2A). A strong acetylation peak was detected for glycyl-tRNA (Figure 2B; Supplementary Table 1), but not for any other tRNAs, indicating that *Mtb* TacT specifically acetylates glycyl-tRNAs.

**Figure 2:**
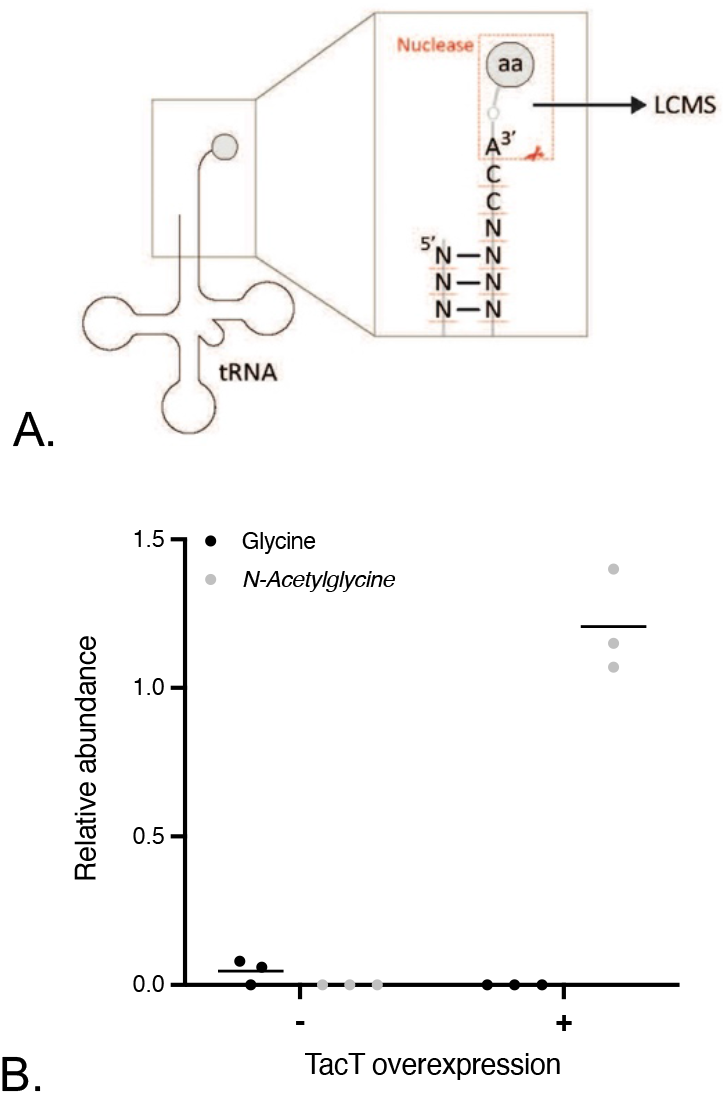
TacT acetylates glycyl-tRNA. *Msmeg* overexpressing *Mtb tacT* was grown to mid-log phase and induced for *tacT* overexpression for 3 hours. Total RNA from triplicate cultures was collected along with an uninduced control for liquid chromatography-mass spectrometry analysis as described. (A) Schematic of nuclease P1 treatment on tRNA samples prior to mass spectrometry. (B) The relative abundance of unacetylated versus N-acetylated glycine is shown as integrated mass spectra peaks normalized to standards.

### Peptidyl tRNA hydrolase (Pth) reverses TacT-induced translation inhibition

Previous work has shown that the enzyme peptidyl tRNA hydrolase (Pth) detoxifies the effects of TacT acetylation by cleaving acetylated amino acids from corrupted tRNAs[14]. To test whether *Mtb* Pth reverses TacT activity, we purified recombinant *Mtb* Pth and added it to our cell-free protein synthesis assay (Supplementary Figure 1). Indeed, purified Pth was sufficient to rescue GFP expression in the presence of active *Mtb* TacT and acetyl coenzyme A (Figure 3B), but had no effects on translation in the presence of catalytically inactive TacT (Figure 3A,B). Thus, *Mtb* Pth also cleaves N-acetylated aminoacyl-tRNA thereby counteracting the effect of the toxin.

**Figure 3:**
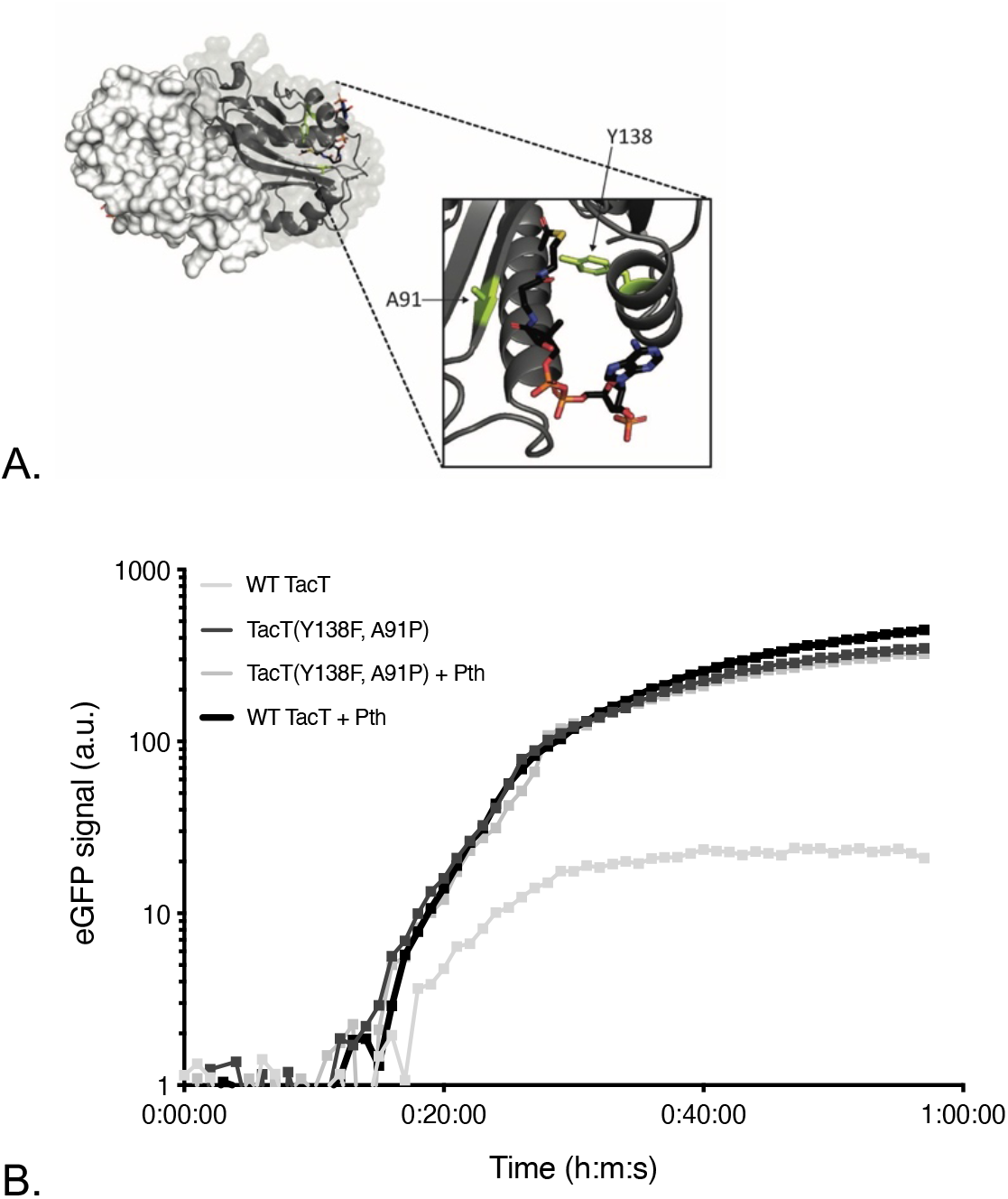
*Mtb* Pth detoxifies TacT. (A) Model of *Mtb* TacT dimer, with one monomer showing mutations for catalytic inactivation (Y138F and A91P; green). Acetyl coenzyme A is shown in the TacT binding pocket and colored by element. (B) Cell-free protein synthesis reactions were set up as described in Figure 1. Inactive TacT was made using an Rv0919 expression construct containing the active side residue mutations Y138F and A91P. Purified *Mtb* peptidyl tRNA hydrolase (Pth) was added where indicated (8uM). In reactions without Pth, an equal volume of storage buffer was added. Protein synthesis was read out as eGFP synthesis and monitored spectrophotometrically at excitation of 488nm and emission of 509nm.

### *tacAT* does not affect *pth* essentiality in *Mtb*

In addition to reversing the effects of TacT-like toxins, Pth’s primary known function is to cleave short peptides from peptidyl-tRNAs that are prematurely released from stalled ribosomes[23, 24]. As with other bacteria, transposon sequencing (TnSeq) data suggest that *pth* is essential in *Mtb*[20]. We built *Mtb pth* transcriptional knockdowns using CRISPR interference (CRISPRi)[25]. Cells induced for *pth* depletion show a marked growth defect, confirming that Pth is required for normal growth (Figure 4). Given TacAT’s connection to this essential enzyme, we assessed whether it contributes to *pth* essentiality in *Mtb*.

**Figure 4:**
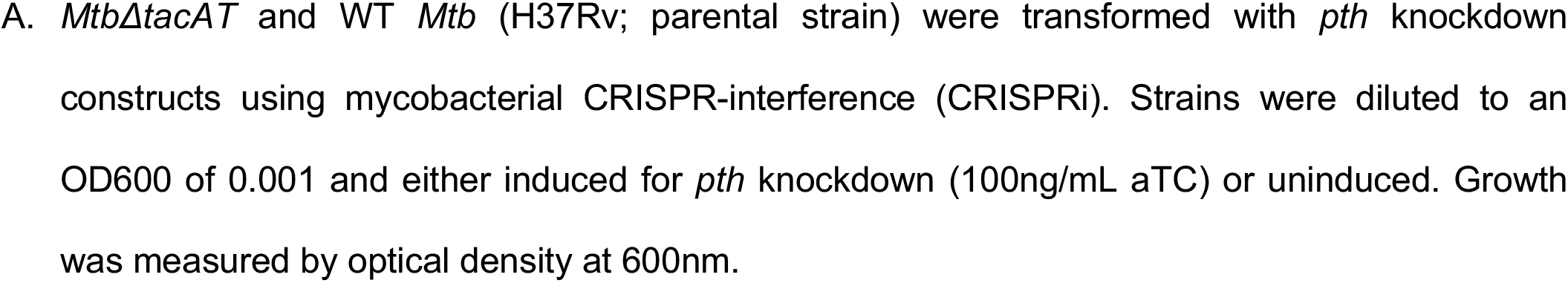
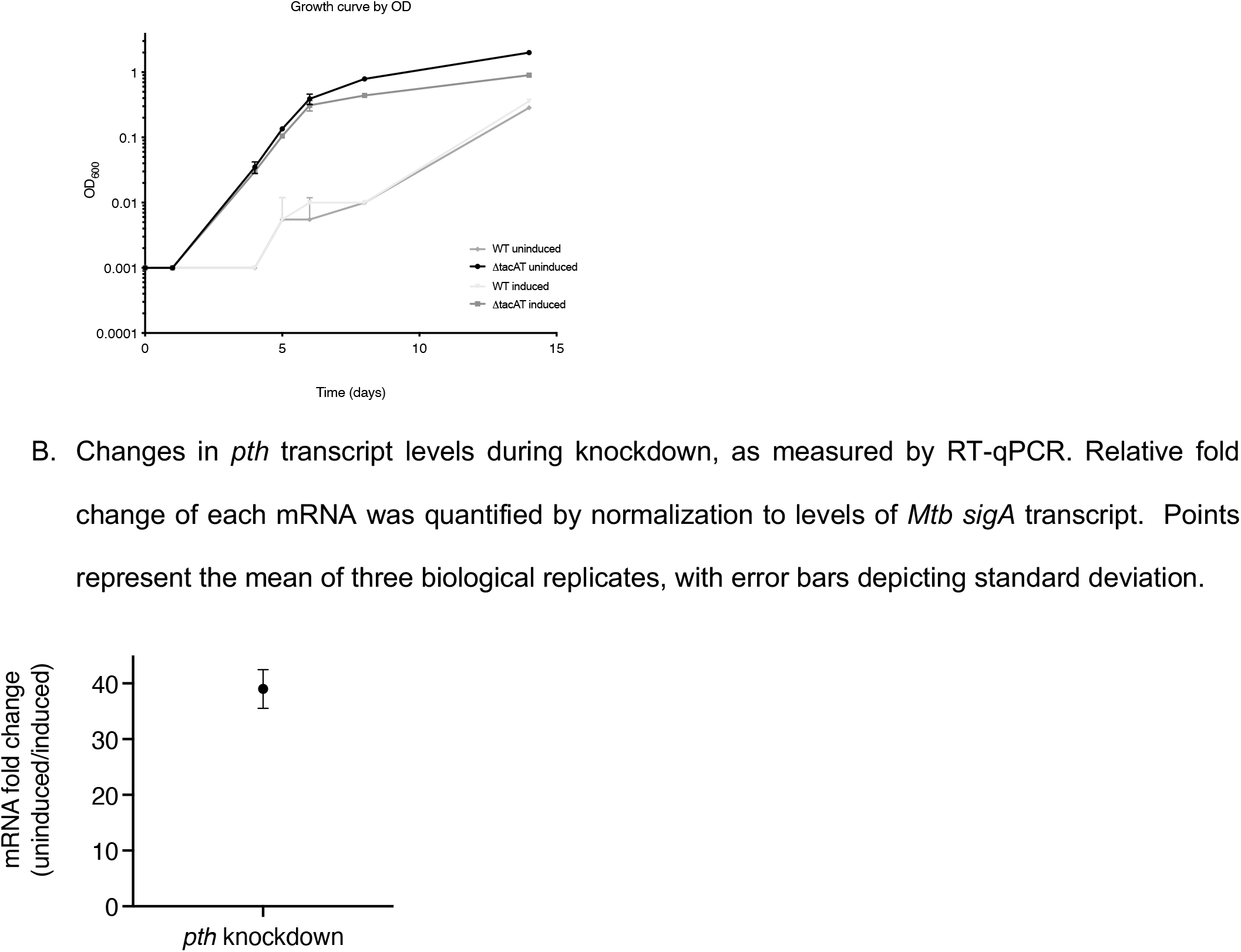
*pth* is still required for normal growth of a *Mtb tacAT* knockout.

Transposon sequencing data indicate that the TacAT operon Rv0918-0919 is nonessential for *Mtb* growth *in vitro*, and we have identified premature stop mutations in this TA system in clinical isolates, suggesting it is also dispensable *in vivo* (Supplementary Table 2)[20]. We built in-frame deletions of the *tacAT* operon in *Mtb* and used CRISPRi to deplete *pth* in this strain. In the absence of *tacAT, pth* knockdowns still failed to grow normally *in vitro*, suggesting that while the *tacAT* operon is dispensable, *pth* essentiality is unaffected by the absence of this TA system (Figure 4). We also performed LCMS on *Mtb* induced or uninduced for *pth* knockdown and were unable to detect glycyl-tRNA acetylation in either strain grown (Supplementary Figure 2). Thus, growth defects of a *pth* knockdown *in vitro* are not a result of the accumulation of acetylated glycyl-tRNAs.

## Discussion

Toxin-antitoxin (TA) systems have been identified in most bacterial genomes and have been implicated in a variety of physiological functions ranging from phage protection and plasmid maintenance to pathogenesis and the general stress response. Interestingly, *Mycobacterium tuberculosis* (*Mtb*) encodes one of the largest repertoires of TA systems in bacteria, yet plasmids are absent from this organism [26]. Furthermore, the role of TA systems in *Mtb* against bacteriophages is still under study [27]. It is tempting to speculate that *Mtb*’s broad TA system toolkit serves as a growth regulator during human infection. However, experimental evidence for this is lacking, largely due to the difficulties of systematically deleting many genes simultaneously in *Mtb* and overlapping mechanisms of action that make it difficult to directly measure the activity of individual toxins. Some studies have examined the roles of individual TA systems in *Mtb* using genetic deletions and overexpression systems, and linked activity of some toxins to pathogenesis[10, 28, 29]. Nonetheless, the level and spectrum of TA system involvement during *Mtb* infection remains unresolved.

Here, we have identified and characterized a TA system in *Mtb* whose mechanism of action is distinct from the other known TA systems in this organism. While most toxins in *Mtb* are ribonucleases, TacT instead blocks growth by acetylating charged tRNAs. This activity can be detected using liquid chromatography-coupled mass spectrometry, making it an appealing TA system to study in its native form. We have shown that *Mtb* TacT acetylates glycyl tRNAs using an overexpression construct but have been unable to detect this modification in wild type *Mtb*. Future work that increases the sensitivity and throughput of LCMS-based or other forms of detection for tRNA acetylation will allow researchers to probe the effects of various physiological conditions on this tRNA modification and identify conditions during which TacT is activated in *Mtb* and in other bacteria containing homologous TA systems.

Recent work using genome sequence from clinical isolates of *Mtb* has shed light on the selective pressures imposed on *Mtb*’s genome during human infection[30, 31]. The essentiality of a gene is correlated with its level of tolerance for nonsynonymous mutations[32]. We have found that the TacAT operon in *Mtb* is dispensable *in vitro*, and clinical genomic data support that this operon is also dispensable *in vivo*, given many nonsynonymous mutations – including premature stop codons – that have accumulated in clinical strains.

The other unique aspect of TacAT is its mechanistic connection to the essential enzyme peptidyl tRNA hydrolase (Pth), which reverses TacT-induced aminoacyl tRNA acetylation. Pth is ubiquitous and thought to be essential across bacteria; in fact, the three critical active site residues His22, Asp95 and Asn116 are universally conserved[33]. Archaea, meanwhile, encode a conserved functional homolog, *pth2*, which does not share significant sequence similarity to bacterial *pth*[33]. Most eukaryotes contain both *pth* and *pth2* genes, though these enzymes are individually nonessential. Interestingly, structural studies have found that mycobacterial Pth is divergent from other bacterial Pth in several regions[34, 35]. Because of its essentiality in bacteria and unique structure in mycobacteria, in addition to the vulnerability of translation rescue systems in *Mtb*, Pth has been proposed as an intriguing drug target in difficult-to-treat organisms like *Mtb* [34, 36, 37]. Understanding the critical functions of Pth is important from a drug development perspective, especially when considering potential sources for antibiotic resistance. For instance, if TacT activity were a significant source for *pth* essentiality in *Mtb*, then inhibitors targeting Pth would lose efficacy in clinical isolates with a disrupted *tacAT* operon. Our work has shown that *Mtb* TacT’s connection to Pth is not the source of *pth* essentiality. This bodes well for studies of Pth as an antibiotic target since mutations inactivating *tacAT* have already been identified in clinical isolates. Finally, biochemical assays to assess Pth inhibitors can exploit the relationship between Pth and TacT using *in vitro* protein synthesis kits by screening for loss of Pth-mediated detoxification.

**Supplementary Figure 1:**
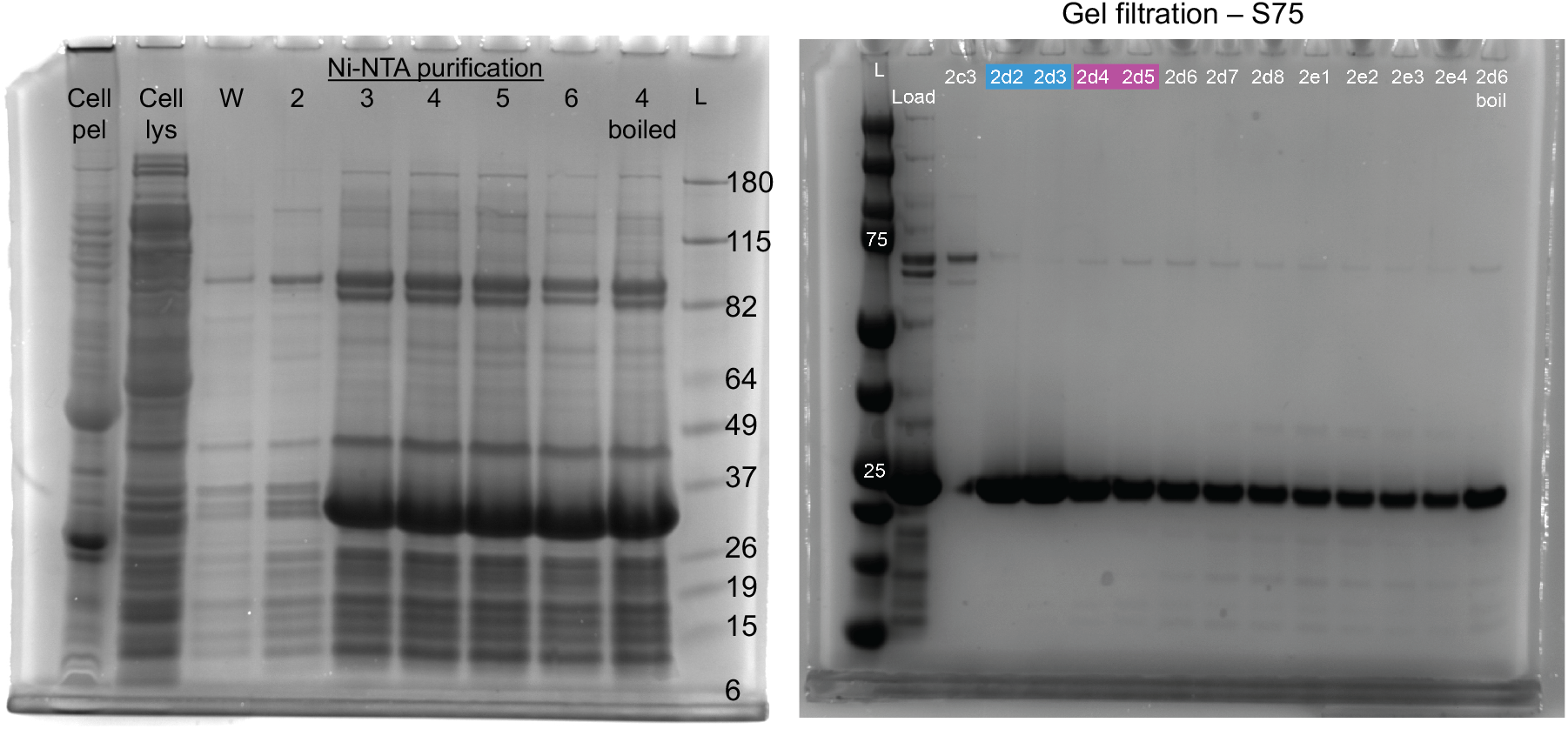
*Mtb* Pth purification gels.

**Supplementary Table 1: Raw LCMS data**.

**Supplementary Table 2: Premature stop mutations in the *tacAT* operon have been detected in clinical *Mtb* isolates (51**,**229 isolates screened)**.

**Supplementary Figure 2:**
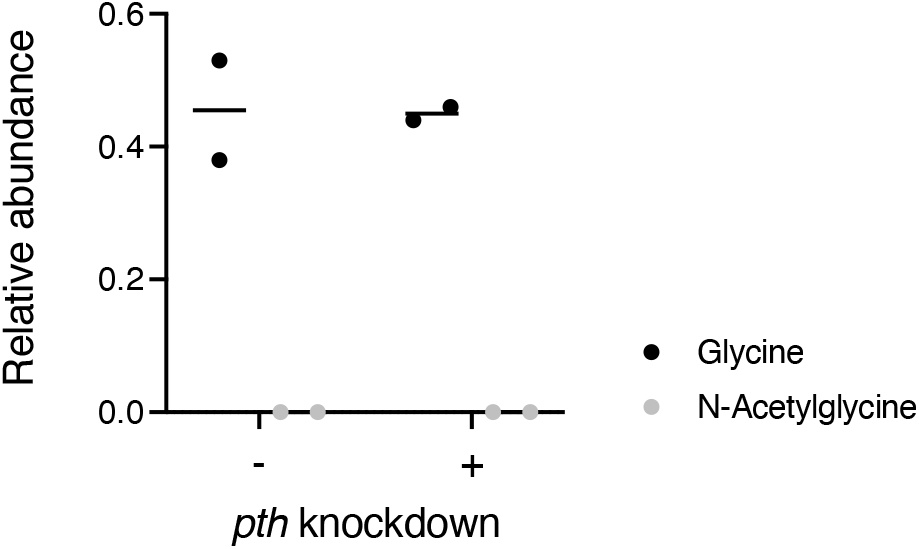
TacT is not active in normal *Mtb* laboratory growth conditions. WT *Mtb* was induced for *pth* depletion and incubated for 4 days. Total RNA from duplicate cultures was collected along with an uninduced control for liquid chromatography-mass spectrometry analysis as described in the Materials and Methods. The relative abundance of unacetylated versus N-acetylated glycine is shown as integrated mass spectra peaks normalized to standards.

**Supplementary Figure 3:**
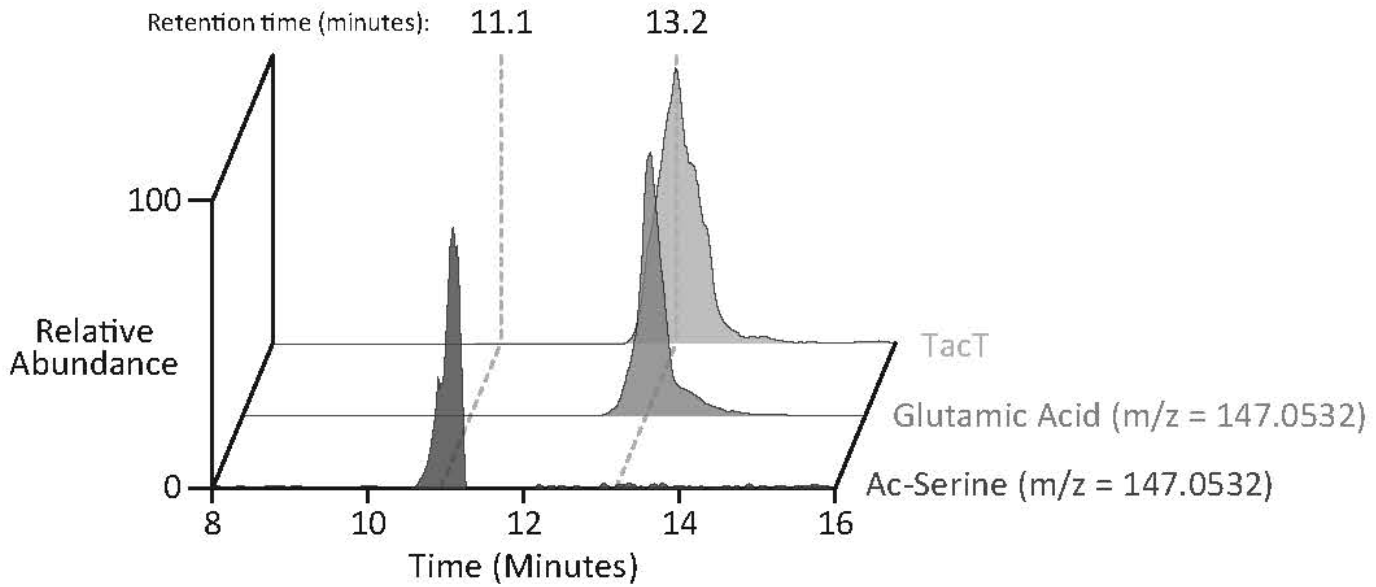
Distinction between acetylated seryl-tRNA and glutamine tRNA. RNA samples treated with Pth were compared to purified standards of each amino acid as described in the Materials and Methods.

## MATERIALS AND METHODS

### Bacterial strains and growth conditions

Mtb and Msmeg strains were grown from frozen stocks into Middlebrook 7H9 medium supplemented with 0.2% glycerol, 0.05% Tween-80, and ADC (5g/L bovine serum albumin, 2g/L dextrose, 3 μg/ml catalase). Cultures were incubated at 37 °C. Antibiotics or inducing agents were used when needed at the following concentrations in both Mtb and Msmeg: kanamycin (25μg/ml), anhydrous tetracycline (aTC; 100ng/mL), hygromycin (50μg/ml), and nourseothricin (20μg/ml). Transformed Mtb and Msmeg strains were plated onto 7H10 agar plates with the appropriate antibiotic(s). Strains were grown to mid log-phase for all experiments unless otherwise specified (OD_600_ 0.4-0.6). *E. coli* strains for cloning or protein purification were grown in LB broth or on LB agar with appropriate antibiotics at the following concentrations: kanamycin (50 μg/ml), zeocin (50μg/ml), and nourseothricin (40 μg/ml). Induction time for *pth* depletion in *Mtb* was 4 days. Induction for *tacT* overexpression in *Msmeg* was 3 hours.

### Bacterial strain construction

Supplementary Table 3 depicts the strains, plasmids, primers, and recombinant DNA used for this study. Plasmids were built by restriction digest of a parental vector and inserts were prepared either by restriction enzyme cloning or Gibson assembly [38] using 40bp overhangs, as specified in Supplementary Table 3. Plasmids were isolated from *E. coli* and confirmed via Sanger sequencing carried out by Genewiz, LLC (Massachusetts, USA).

#### Deletion mutants

The knockout strain Δ*tacAT*::zeo (zeocin) was built using double-stranded recombineering in the parental *Mtb* strain H37Rv. A linear dsDNA fragment was constructed using stitch PCR with the primers listed in Supplementary Table 3 which consisted of a 500bp region upstream of the *tacAT* operon (Rv0918-019), 500bp downstream region, and a *lox-*zeo*-lox* fragment. This cassette was transformed into an H37Rv recombineering strain as described [39] and plated on 7H10 + zeocin plates.

#### *tacAT, tacT* alleles

Plasmid FT2, used for inducible *tact* overexpression in *Msmeg*, was generated using a parental vector (CT16) that integrates into the L5 mycobacterial phage site. This plasmid also encodes for kanamycin resistance and contains both the tet promoter (directly upstream of *tacT*) and the tet repressor. CT16 was digested with ClaI and XbaI (New England Biolabs). *tacT* (Rv0919) was PCR-amplified and ligated into the plasmid using restriction cloning. Plasmid FT3, used for *tacAT* overexpression, was generated by placing *tacAT* together under the constitutive UV15 promoter in a parental vector (CT250) which was digested with NdeI and HindIII (New England Biolabs). The *tacAT* operon Rv0918-0919 was ligated to the plasmid using Gibson cloning.

#### Pth-knockdown constructs

Transcriptional knockdown of *pth* was accomplished using mycobacterial CRISPRi-interference (CRISPRi). Knockdown constructs were built as previously described [25] by annealing oligos for *pth* and ligating them into a linearized BsmBI-digested plasmid (CT 296; gift of Jeremy Rock) that contains mycobacterial CRISPRi. The knockdown vector FT110 was transformed in both H37Rv wild type (WT) and ΔtacAT::ZeoR.

### Purification of *Mtb* Pth

*Mtb pth* (Rv1014c) was cloned with a C-terminal 6x His-tag and expressed from pET28a in BL21-CodonPlus (DE3)-RP *E. coli* under conditions similar to those previously described[40]. 1 L of log-phase culture (OD600 ~ 0.7) was induced with 1 mM isopropyl β-D-1-thiogalactopyranoside (IPTG) for 4 hours at 37°C. Cells were harvested at 6,000g for 15 minutes, and the resulting pellet was frozen at −80°C. The pellet was thawed with a stir-bar at 4°C in lysis buffer containing 50 mM Tris HCl pH 7.5, 300 mM NaCl, 10% glycerol, a pinch of DNase powder, 1 tablet ecomplete EDTA-free protease inhibitor and 2 mM 2-mercaptoethanol (BME), and cells were lysed using a French press. Lysate was clarified by spinning at 30,000g for 30 minutes and brought up to 20 mM imidazole pH 7.5. His-tagged Pth was then extracted via batch binding 2.5 mL equilibrated Ni-NTA beads incubated with lysate for 1 hour at 4°C. Beads were collected and washed with 20 mL lysis buffer containing 2 mM BME. A second wash included 20 mL lysis buffer with 20 mM imidazole and 2 mM BME, followed by 5 mL of lysis buffer with 30 mM imidazole, and a final wash with 5 mL lysis buffer containing 40 mM imidazole and 2 mM BME. Samples were eluted with lysis buffer containing 200 mM imidazole pH 7.5 in 750 μL fractions and analyzed via SDS-PAGE (Supplementary Figure 1, left). The cleanest elution fractions (4-6 and 7-9) were desalted into lysis buffer containing BME, concentrated with a 10 KDa MWCO amicon ultra 4 spin column to 1 mL and further purified by FPLC via gel filtration chromatography with a Superdex 75 Increase 10/300 GL column in buffer containing (25 mM Tris-HCl pH 7.5, 150 mM NaCl, 2 mM BME). Fractions were analyzed via SDS-PAGE (Supplementary Figure 1, right) and fractions 2d2-2d3 (at an elution volume ~13 mL) were pooled and brought up to 5% glycerol with 2 mM fresh BME. Nanodrop readings suggested that other fractions containing what appeared to be pure Pth were contaminated by unknown nucleic acid species. Nucleic acid-free protein (fractions 2d2-2d3) was aliquoted into 10 uL aliquots, flash frozen with liquid nitrogen, and stored at −80°C. Pth protein concentration was calculated using a Coomassie Plus (Bradford) Assay (Pierce).

### *In vitro* translation

To assess the effect of TacT on translation, *in vitro* translation reactions were prepared with purified *tacT* DNA (WT, Y138F, or Y138F/A91P), 2nM acetyl coenzyme A, and purified Pth. A master mix of purified eGFP DNA (200ng per reaction), *tact* DNA (180ng per reaction) and PURExpress (New England Biolabs) components were prepared in triplicate reactions with 8μM of Pth and 2nM acetyl coenzyme A. When no Pth was added, an equal volume of storage buffer was used in place of protein. When no acetyl coenzyme A was added, an equal volume of water was added. Reactions were carried out in 12μL in a black Co-star 384-well plate for 2 hours at 37°C, and eGFP fluorescence (excitation = 488 nm and emission = 509 nm) was measured over time on a SpectraMax M2 microplate reader.

### mRNA quantification

10mL *Mtb* cultures were harvested at 4,000rpm for 10 minutes and pellets were resuspended in 1mL TriZol reagent (ThermoFisher Scientific). Samples were lysed by bead beating. Purified DNase-treated RNA was used as template for cDNA synthesis, following manufacturer’s instructions with Superscript IV (Life Technologies). RNA was removed using RNase A (ThermoFisher Scientific) and cDNA cleaned up by column purification (Zymo Research). qPCR was performed using iTaq Universal SYBR Green Supermix (BioRad). mRNA fold-change was calculated using the ΔΔCt method, where *pth* transcript level was normalized by *sigA* level in each condition.

### Liquid chromatography-coupled mass spectrometry

Purified RNA (15 – 50 μg) was incubated with 1U Nuclease P1 in 10mM ammonium acetate for 30 minutes at 25 °C (Figure 2 and Supplementary Figure 2), or with 25 µg purified Pth in buffer (10 mM Tris acetate, 10mM magnesium acetate, 20mM ammonium acetate pH 8.) for 1 hour at 37 °C (Supplementary Figure X). Processed RNA samples were diluted 1:3 with acetonitrile + 0.2% v/v acetic acid, centrifuged for 10 minutes at 21,000 g, room temperature to remove any precipitate, and transferred to glass microvials. Samples were analysed on a Thermo Ultimate 3000 LC coupled with a Q-Exactive Plus mass spectrometer in both positive and negative ion modes. Five microliters of each sample were injected on a Zic-pHILIC Column (150×2.1 mm, 5 micron particles, EMD Millipore). The mobile phases are (A) 20 mM ammonium carbonate in 0.1 % ammonium hydroxide and (B) acetonitrile 97% in water. The gradient conditions were as follows: 100% B at 0 min, 40% B at 20 min, 0% B at 30 min for 5 min, then back to 100% B in 5 min, followed by 10 min of re-equilibration. A constant flow rate of 0.200 L/minute was used. The mass spectrometer was calibrated immediately prior to use. Data were analyzed using Thermo Xcalibur 3.0 with ICIS automated peak integration (Default settings: Smoothing Points = 9; Baseline Window = 40; Area Noise Factor = 2; Peak Noise Factor = 10) followed by manual data curation. To distinguish the isobaric molecules N-acetylserine and glutamic acid using LCMS, RNA samples treated with Pth were compared to purified standards of each amino acid. These data indicate that glutamic acid, and not N-acetylserine, contributes the entirety of the MS signal detected for molecules with a mass of 147.0532 (Supplementary Figure 3).

### Whole genome sequencing analysis of clinical isolates

Whole genome sequences of 55778 Mtb isolates were obtained from 211 BioProjects under the following accession codes: ERP001037, ERP002611, ERP008770, PRJDB10607, PRJDB3875, PRJDB6149, PRJDB7006, PRJDB8544, PRJDB8553, PRJEB10385, PRJEB10533, PRJEB10577, PRJEB10950, PRJEB11460, PRJEB11653, PRJEB11778, PRJEB12011, PRJEB12179, PRJEB12184, PRJEB12764, PRJEB13325, PRJEB13764, PRJEB13960, PRJEB14199, PRJEB15076, PRJEB15382, PRJEB15857, PRJEB18529, PRJEB20214, PRJEB21685, PRJEB21888, PRJEB21922, PRJEB23245, PRJEB23495, PRJEB2358, PRJEB23648, PRJEB23664, PRJEB23996, PRJEB24463, PRJEB25506, PRJEB25543, PRJEB25592, PRJEB25814, PRJEB25968, PRJEB25971, PRJEB25972, PRJEB25991, PRJEB25997, PRJEB25998, PRJEB25999, PRJEB26000, PRJEB26001, PRJEB26002, PRJEB27244, PRJEB27354, PRJEB27366, PRJEB27446, PRJEB27847, PRJEB2794, PRJEB28497, PRJEB28842, PRJEB29199, PRJEB29276, PRJEB29408, PRJEB29435, PRJEB29446, PRJEB29604, PRJEB30463, PRJEB30782, PRJEB30933, PRJEB31023, PRJEB31905, PRJEB32037, PRJEB32234, PRJEB32341, PRJEB32589, PRJEB32684, PRJEB32773, PRJEB33896, PRJEB35201, PRJEB39699, PRJEB40777, PRJEB5162, PRJEB5280, PRJEB5899, PRJEB5925, PRJEB6273, PRJEB6717, PRJEB6945, PRJEB7056, PRJEB7281, PRJEB7669, PRJEB7727, PRJEB7798, PRJEB8311, PRJEB8432, PRJEB8689, PRJEB9003, PRJEB9201, PRJEB9206, PRJEB9308, PRJEB9545, PRJEB9680, PRJEB9709, PRJEB9976, PRJNA200335, PRJNA217391, PRJNA219826, PRJNA220218, PRJNA229360, PRJNA233386, PRJNA235852, PRJNA237443, PRJNA244659, PRJNA254678, PRJNA259657, PRJNA268900, PRJNA270137, PRJNA282721, PRJNA287858, PRJNA295328, PRJNA300846, PRJNA302362, PRJNA305488, PRJNA306588, PRJNA308536, PRJNA318002, PRJNA352769, PRJNA353873, PRJNA354716, PRJNA355614, PRJNA356104, PRJNA361483, PRJNA369219, PRJNA376471, PRJNA377769, PRJNA379070, PRJNA384604, PRJNA384765, PRJNA384815, PRJNA385247, PRJNA388806, PRJNA390065, PRJNA390291, PRJNA390471, PRJNA393378, PRJNA393923, PRJNA393924, PRJNA401368, PRJNA401515, PRJNA407704, PRJNA413593, PRJNA414758, PRJNA419964, PRJNA421323, PRJNA421446, PRJNA428596, PRJNA429460, PRJNA430531, PRJNA431049, PRJNA436223, PRJNA436997, PRJNA438921, PRJNA448595, PRJNA453687, PRJNA454477, PRJNA475130, PRJNA475771, PRJNA480117, PRJNA480888, PRJNA481625, PRJNA481638, PRJNA482095, PRJNA482716, PRJNA482865, PRJNA486713, PRJNA488343, PRJNA488426, PRJNA492975, PRJNA506272, PRJNA509547, PRJNA512266, PRJNA522942, PRJNA523164, PRJNA523499, PRJNA524863, PRJNA526078, PRJNA528965, PRJNA533314, PRJNA540911, PRJNA549270, PRJNA559678, PRJNA566379, PRJNA573497, PRJNA578162, PRJNA586859, PRJNA587747, PRJNA589048, PRJNA591498, PRJNA595834, PRJNA598949, PRJNA598981, PRJNA608715, PRJNA632617, PRJNA663350, PRJNA678116, PRJNA679443, PRJNA683067, PRJNA684613, PRJNA688213, SRA065095. The Sickle tool was used for trimming whole-genome sequencing data[41]. Sequencing reads with Phred base quality scores above 20 and read lengths longer than 30 were kept for analysis. The inferred ancestral genome of the most recent common ancestor of the MTBC was used as the reference template for read mapping[42]. Sequencing reads were mapped to the reference genome using Bowtie 2 (version 2.2.9) [43]. SAMtools (v1.3.1) was used for SNP calling with mapping quality greater than 30. Fixed mutations (frequency ≥ 75%) were identified using VarScan (v2.3.9) with at least 10 supporting reads and the strand bias filter option on. SNPs in repetitive regions of the genome (e.g., PPE/PE-PGRS family genes, phage sequences, insertion or mobile genetic elements) were excluded [44, 45].

## Supporting information

STOPmutations

## Data availability statement

The data that support these findings are available from the corresponding author upon reasonable request.

## Author contributions

Conceptualization: F.G.T., J.T.P.S., E.J.R.; Methodology: F.G.T, A.M.J.H., J.T.P.S., C.L.D., K.M.; Investigation: F.G.T., A.M. J.H., J.T.P.S., C.L.D., K.M.; Data Curation: F.G.T., A.M. J.H., Q.L. Writing – Original Draft: F.G.T., E.J.R.; Writing – Review & Editing: F.G.T., A.M. J.H., J.T.P.S., C.L.D., K.M., Q.L., S.M.F., S.H., E.J.R.

